# Combining *in vivo* and *in vitro* approaches to better understand host-pathogen interactions

**DOI:** 10.1101/2024.04.19.590205

**Authors:** Robert Holdbrook, Catherine E. Reavey, Joanna L. Randall, Awawing A. Andongma, Yamini Tummala, Stephen J. Simpson, Judith A. Smith, Sheena C. Cotter, Kenneth Wilson

**Affiliations:** Lancaster Environment Centre, Lancaster University, Lancaster LA1 4YQ, UK; Charles Perkins Centre, The University of Sydney, NSW 2006, Australia; School of Forensic and Applied Sciences, University of Central Lancashire, Preston, Lancashire, PR1 2HE, UK; School of Life Sciences, University of Lincoln, Brayford Pool, Lincoln LN6 7TS, UK

**Keywords:** Nutritional geometry, immunity, *Spodoptera*, *Xenorhabdus*, pathogenic bacteria

## Abstract

Nutrition often shapes the outcome of host-parasite interactions, however understanding the mechanisms by which this occurs is confounded by the intimate nature of the association and by the fact that the host and parasite may compete for the same limiting nutrients. One way of disentangling this interaction is to combine *in vivo* and *in vitro* approaches. Here, we explore the role of host nutrition in determining the outcome of infections using a model insect-bacterium system: the cotton leafworm *Spodoptera littoralis* and the blood-borne bacterium *Xenorhabdus nematophila*.

*Spodoptera littoralis* larvae were reared on one of a series of 20 chemically-defined diets ranging in their protein: carbohydrate (P:C) ratio and caloric density. They were then challenged with either a fixed dose of *X. nematophila* cells (live or dead) or were sham-injected. Survivorship of larvae challenged with live bacterial cells was strongly dependent on the protein levels of the diet, with mortality being highest on low-protein diets. This trend was reflected in the bacterial growth rate *in vivo*, which peaked in larvae fed low-protein diets.

To determine whether *in vivo* bacterial growth rates were driven by blood nutrients, rather than an enhanced host immune response, we generated 20 synthetic haemolymphs (‘nutribloods’) that mimicked the nutritional content of host blood. Bacterial growth rate in the nutribloods was also negatively impacted by their protein content suggesting that nutrient availability and not host immunity was driving the interaction. By comparing standardized bacterial growth rates *in vivo* and *in vitro*, we conclude that the outcome of this host-parasite interaction is largely driven by ‘bottom-up’ effects of nutrients on bacterial growth, rather than by ‘top-down’ effects of nutrients on host-mediated immune responses. The outcome of host-parasite interactions is typically assumed to be strongly determined by the host immune response. The direct effects of nutrition have been underexplored and may have broad consequences for host-parasite interactions across taxa.

## INTRODUCTION

Nutritional immunology explores the role of nutrient availability in the delicate balance between hosts and their parasites (Ponton et al., 2011, 2013, 2023). Much of this research has focused on the effects of nutrients on host immune function and/or on the outcome of an infection, with host fitness often being positively correlated with elevated levels of host immunity (Ponton et al., 2023; Schmid-Hempel, 2021; Wilson et al., 2019). What is often overlooked in these studies is the effects of host nutrition on parasite establishment and proliferation. This is an important knowledge gap because parasites are usually dependent on their hosts for nutritional resources and both the host and its parasite may compete for the same limiting nutrients. Thus, the outcome of an infection may be determined primarily by the effects of host nutrition on ‘top down’ (e.g. immunological) processes directed at the parasite, ‘bottom up’ effects of specific nutrients on parasite population growth, or a combination of the two (Griffiths et al., 2015; Haydon et al., 2003; Metcalf et al., 2011; Mideo & Reece, 2012; Moore et al., 2018; Ramiro et al., 2016). Disentangling the relative importance of top-down and bottom-up regulation of parasites is difficult due to the intimate nature of the association, but it is possible by combining *in vivo* and *in vitro* approaches in a tractable system.

Here, we use as a model system for teasing apart a host-parasite interaction in a generalist caterpillar host, the cotton leafworm *Spodoptera littoralis*, and its parasite *Xenorhabdus nematophilus*, an extra-cellular gram-negative bacterium. The bacterium has a mutualistic association with the entomopathogenic nematode *Steinernema carpocapsae*, which vectors the bacterium into insect hosts via the cuticle or orifices such as the mouth and anus. Importantly, *X. nematophila* is also able to kill its insect host without the nematode when it is injected directly into the insect haemocoel, making it a tractable system (Wilson et al., 2020). *S. littoralis* is a useful host because it is relatively easy to culture in the laboratory, it is a generalist feeder, and large amounts of haemolmph can be extracted from a single individual, allowing multiple blood tests to be undertaken.

Nutritional geometry (NG) is a state-space nutritional modelling approach that is aimed at determining the effects of multiple nutrients on an organism’s behaviour and fitness (Simpson & Raubenheimer, 1995). A previous study developed twenty chemically-defined diets systematically varying in their concentration and balance of two key macronutrients, proteins and carbohydrates (Cotter et al., 2011). These twenty diets reflect the variation in these macronutrients that *S. littoralis* would naturally encounter in its environment (**Figure S1**). Using these diets, we showed that some aspects of *S. littoralis* immune responses are heightened in a high-protein environment (Cotter et al., 2011, 2019; Lee et al., 2006), demonstrating the potential for top-down (immunological) effects on bacterial growth. In a more recent study (Holdbrook, Andongma, et al., 2024) we used this approach to establish the effects of host nutrition on the insect haemolymph (blood) nutrient pool for insect feeding on the same 20 chemically-defined diets. This established that whilst carbohydrates in the haemolymph are generally tightly regulated, haemolymph protein concentration tends to increase with the amount of protein eaten (Holdbrook, Andongma, et al., 2024). In a subsequent study, we used this detailed analysis of haemolymph nutrients for the 20 diets to generate 20 synthetic haemolymphs (‘*nutribloods’*) that mimicked the nutritional profile of the real haemolymphs of caterpillars fed the 20 chemically-defined diets (Holdbrook, Randall, et al., 2024). This revealed that the *in vitro* growth of *X. nematophila* (in the absence of host immune defences) increased with the amount of carbohydrate in the nutriblood and decreased with the amount of protein, suggesting potential bottom-up effects of nutrition on bacterial performance.

Here, we combine these *in vitro* nutriblood results with *in vivo* bacterial dynamics to tease apart the relative importance of top-down and bottom-up effects of host nutrition in determining the outcome of infections by *X. nematophila* in *S. littoralis*. The work builds on our earlier study that explored this interaction using just six chemically-defined diets covering a limited range of nutrient space (Wilson et al., 2020). In the present study, we combine *in vitro* and *in vivo* experiments using 20 chemically-defined diets to test the robustness of this finding and to statistically compare bacterial growth in the two settings.

## METHODS

### Cultures

#### Insect culture

The *Spodoptera littoralis* culture was founded in 2002 from eggs collected from Egypt. It was maintained using single-pair matings of over 150 pairs per generation to reduce in-breeding. For experiments, larvae were collected in the 2^nd^ instar and reared singly on a semi-artificial wheatgerm-based diet until the start of the final instar (6^th^). Larvae were kept in 25 mL polypots at 27 °C under a 12:12 light: dark regime.

#### Bacterial culture

Bacteria were originally supplied by the laboratory of Givaudan and colleagues (Montpellier University, France; *X. nematophila* F1D3 GFP labelled). It was maintained on nutrient agar at 4°C and stored in liquid culture at −80°C (1:1 nutrient broth culture:glycerol). Bacteria were used to infect 6^th^ instar *S. littoralis* larvae to maintain virulence and single colonies grown from haemolymph-smeared NBTA agar plates (Sicard et al., 2004) were then grown in sterile nutrient broth for 24 h at 28 °C shaking at 150 rpm. Stocks were made by mixing 500 µL of liquid culture with glycerol at a 1:1 ratio and stored again at −80 °C. Prior to experiments, bacteria were revived from the frozen stores: 100 µl of frozen rpm.

### *In vivo* experiments

#### Bacterial culture

The methods for the *in vivo* experiments are based on those of Wilson et al (2020). In brief, on the day of bacterial challenge, the bacterial stock was sub-cultured in nutrient broth and placed in a shaker-incubator for *c*. 4 h to ensure that the bacteria were in log phase. Following this, the concentration of bacterial cells was quantified using a fluorescence microscope in a serial dilution of nutrient broth using a haemocytometer with improved Neubauer ruling. The remaining culture was further diluted with nutrient broth to the appropriate concentration required for the bacterial challenge. Half of the culture was then autoclaved to use as a ‘dead-bacteria’ control group.

#### Experimental design

Four hundred larvae were reared to the start of 6th instar on a semi-artificial wheat germ-based diet. Within 24 h of moulting into the final instar, the larvae were divided into 20 groups (n=20 larvae) and placed singly onto one of twenty diets differing in dietary attributes (Table S1). Between 1.8 and 2.1 g of the chemically-defined diets were placed in 90 mm diameter Petri dishes and the larvae were housed in this manner throughout the experiment (with the diet replaced every 24 h). Within each diet, 10 caterpillars were allocated to a ‘live bacteria challenged’ group (henceforth live-infected), 5 caterpillars were assigned to a ‘dead bacteria challenged’ group (henceforth dead-infected) and 5 caterpillars were allocated to a ‘sham challenged’ group (henceforth sham-infected). For the live-infected caterpillars, the bacterial dose used was 1,272 *X. nematophila* cells/mL nutrient broth. This dose was established from pilot experiments to determine the LD_50_ (unpublished data). The same dose was used for the dead-infected challenge, albeit the challenge would consist of cell debris as a result of autoclaving. The sham-infected caterpillars were injected with autoclaved nutrient broth only.

Following 24 h on the assigned diets, each of the 400 caterpillars was injected with the appropriate treatment; 5 µL of live *X. nematophila*, 5 µL of heat-killed *X. nematophila* or 5 µL of autoclaved nutrient broth. Injections were carried out using a Hamilton Syringe in a micro-injector. The syringe was sterilized in ethanol before each injection and the challenge was applied to the left proleg nearest to the head. Time of injection was recorded due to the need to control for the length of time between injection of the first and last individuals (4.5 h).

After challenge, haemolymph samples were obtained from all caterpillars at roughly 20 h post-infection. Samples were obtained by piercing the cuticle next to the first proleg near the head with a sterile needle and allowing released haemolymph to bleed directly into an Eppendorf tube. Haemolymph samples from all the live-injected caterpillars were plated out to determine bacterial growth (n = 200). One of each of the 5 caterpillars for both the dead-infected and sham-infected caterpillars within each dietary treatment were plated out to ensure no bacterial contamination had occurred (n = 40). Immediately after obtaining the haemolymph, the relevant samples were diluted in pH 7.4 phosphate buffered saline (PBS; 10 µL of haemolymph placed in 90 µL of PBS and so on through the dilution series) down to 10^-7^ at intervals of 10^-1^. The dilution series was plated onto NBTL agar plates (20 µl per 1/4 agar plate) containing bromothymol blue and triphenyltetrazolium chloride and incubated at 28°C. *Xenorhabdus* colonies appear deep blue on these NBTL agar plates, whereas most other bacterial species appear yellow or red, allowing contaminants to be identified. Although most of the colonies were visible at 24 h, there were some slow growing colonies that were not visible until 48 h. Following the incubation period at 28°C, the CFUs were counted for each sample, and then the CFU/ml haemolymph were determined based on the dilution factor at which colonies could be reliably counted.

Fresh diet was provided in clean 90 mm diameter Petri dishes every 24 h up to 72 h (48 h post infection). Ninety-six hours after moulting into 6th instar, the larvae had either pupated or were placed in pots of semi-artificial diet until death or pupation. All caterpillars were monitored for death throughout the day of sampling and every day after until pupation or death.

### *In vitro* growth experiments

#### Synthetic haemolymphs: nutribloods

The full methods for generating the 20 nutribloods is outlined in Holdbrook et al. (2024). In brief, the design was based on the nutritional composition of *S. littoralis* fed on the 20 chemically-defined diets using a combination of (ultra) high performance liquid chromatography ((u)HPLC) and spectroscopic methods, using Grace’s insect medium (Sigma Aldrich G8142) as a source of minerals and vitamins (Tables 1–3 in Holdbrook, Randall, et al., 2024).

**Table 1.**
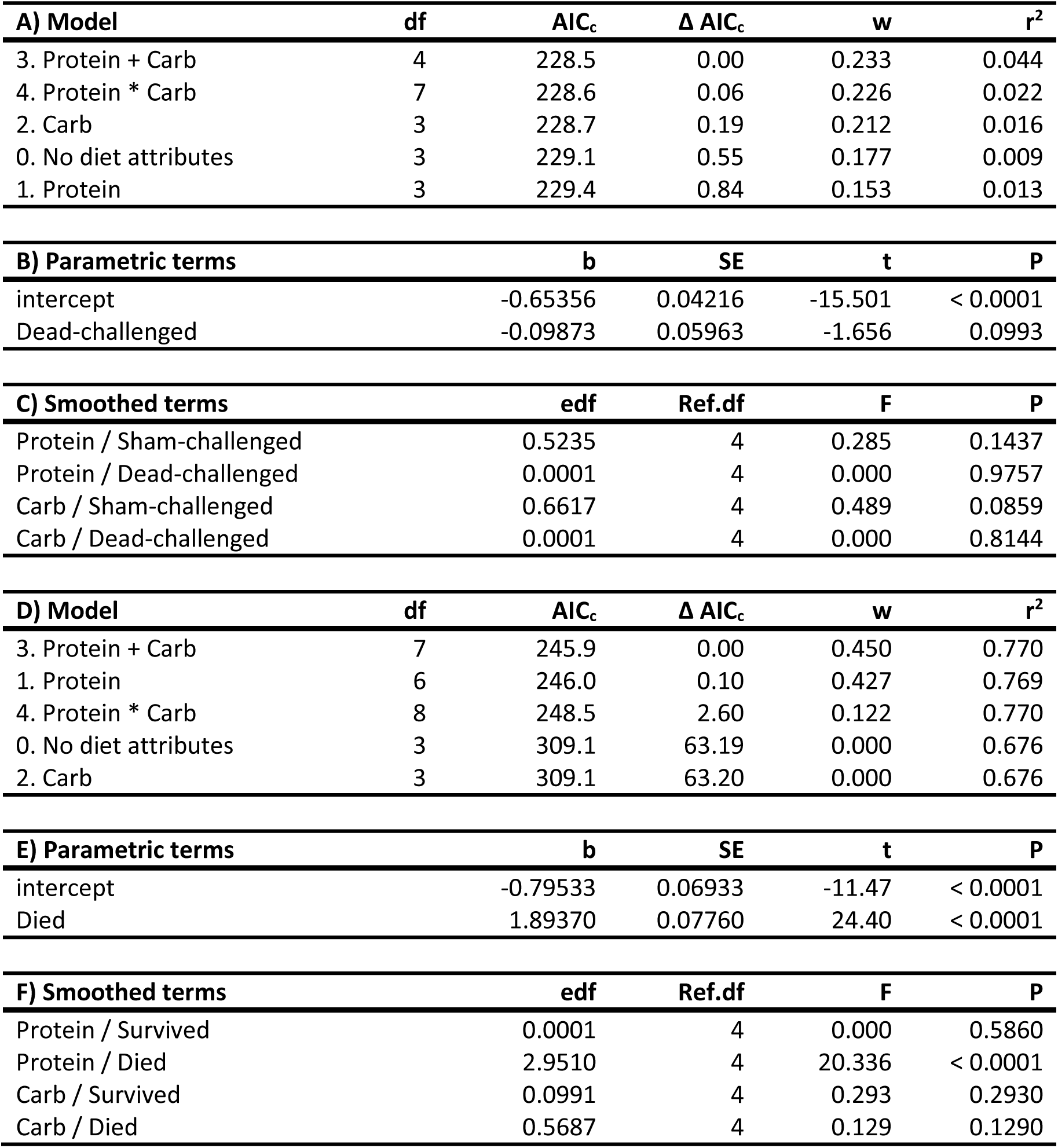
GAMs explaining speed of death in relation to the nutritional attributes of the host diet for larvae in the two control groups (A-C), and for larvae in the live-bacteria challenged group only (D-F). **A,D)** Table of candidate models. df = degrees of freedom, AICc = corrected Akaike Information Criteria values; Δ AICc = difference in AICc values between the best model (lowest AICc) and the given model; w = Akaike weights; r2 = pseudo-r2 for the model. The five alternative Diet attributes listed in the first column are described in Table S1. The dependent variable in these models is the speed of death (1 / time taken to die, h), standardised to have a mean of zero and a standard deviation of unity for all larvae (see Methods). **B,E)** Parametric terms and **C,F)** Smoothed terms of the top model (model 3).

| **A) Model** | **df** | **AIC<sub>c</sub>** | **$\Delta$ AIC<sub>c</sub>** | **w** | **r<sup>2</sup>** |
| --- | --- | --- | --- | --- | --- |
| 3. Protein + Carb | 4 | 228.5 | 0.00 | 0.233 | 0.044 |
| 4. Protein * Carb | 7 | 228.6 | 0.06 | 0.226 | 0.022 |
| 2. Carb | 3 | 228.7 | 0.19 | 0.212 | 0.016 |
| 0. No diet attributes | 3 | 229.1 | 0.55 | 0.177 | 0.009 |
| 1. Protein | 3 | 229.4 | 0.84 | 0.153 | 0.013 |

| **B) Parametric terms** | **b** | **SE** | **t** | **P** |
| --- | --- | --- | --- | --- |
| intercept | -0.65356 | 0.04216 | -15.501 | < 0.0001 |
| Dead-challenged | -0.09873 | 0.05963 | -1.656 | 0.0993 |

| **C) Smoothed terms** | **edf** | **Ref.df** | **F** | **P** |
| --- | --- | --- | --- | --- |
| Protein / Sham-challenged | 0.5235 | 4 | 0.285 | 0.1437 |
| Protein / Dead-challenged | 0.0001 | 4 | 0.000 | 0.9757 |
| Carb / Sham-challenged | 0.6617 | 4 | 0.489 | 0.0859 |
| Carb / Dead-challenged | 0.0001 | 4 | 0.000 | 0.8144 |

| **D) Model** | **df** | **AIC<sub>c</sub>** | **$\Delta$ AIC<sub>c</sub>** | **w** | **r<sup>2</sup>** |
| --- | --- | --- | --- | --- | --- |
| 3. Protein + Carb | 7 | 245.9 | 0.00 | 0.450 | 0.770 |
| 1. Protein | 6 | 246.0 | 0.10 | 0.427 | 0.769 |
| 4. Protein * Carb | 8 | 248.5 | 2.60 | 0.122 | 0.770 |
| 0. No diet attributes | 3 | 309.1 | 63.19 | 0.000 | 0.676 |
| 2. Carb | 3 | 309.1 | 63.20 | 0.000 | 0.676 |

| **E) Parametric terms** | **b** | **SE** | **t** | **P** |
| --- | --- | --- | --- | --- |
| intercept | -0.79533 | 0.06933 | -11.47 | < 0.0001 |
| Died | 1.89370 | 0.07760 | 24.40 | < 0.0001 |

| **F) Smoothed terms** | **edf** | **Ref.df** | **F** | **P** |
| --- | --- | --- | --- | --- |
| Protein / Survived | 0.0001 | 4 | 0.000 | 0.5860 |
| Protein / Died | 2.9510 | 4 | 20.336 | < 0.0001 |
| Carb / Survived | 0.0991 | 4 | 0.293 | 0.2930 |
| Carb / Died | 0.5687 | 4 | 0.129 | 0.1290 |

#### Preparation of bacteria

The methods for preparing the bacteria are outlined in full in Holdbrook et al. (2024). In brief, bacteria were revived from frozen liquid stores and sub-cultured in nutrient broth before being incubated for 4 h to reach log phase of growth. The bacterial cells were washed in PBS to avoid the transfer of nutrients from nutrient broth into the growth media, following Crawford et al (2012). A 1 mL sample was used to generate a dilution series in GIM-saline from which the total cell count was determined using a haemocytometer with improved Neubauer ruling. The remaining culture was then diluted in (Sprouffske, 2020).

#### Bacterial growth assays

Bacterial cell growth was quantified using a SpectraMax Plus microtiter plate reader (Molecular Devices) with SoftMax Pro software (Holdbrook et al. manuscript). Each plate contained 180 µL of one of the 20 nutribloods in quadruplets. The turbidity at 600 nm was determined every 10 min for 30 h and the plate was shaken for 30 s before each measurement (Holdbrook, Randall, et al., 2024).

### Data analysis

All data analyses was performed using the R statistical software (v4.3.0; R Core Team, 2014). The *in vitro* bacterial growth kinetics were quantified using the *Growthcurver* package (v0.3.1 (Sprouffske, 2020), with the maximum optical density (OD) at 600 nm. Insect survivorship post-challenge until death (in the larval, pupal or moth stage) was analysed by fitting a parametric survival regression model, using the *survreg* function in the survival package (v3.5.5; (Therneau, 2023)). Speed of death was quantified as 1/time to death (h). Both *in vivo* and *in vitro* data were analysed using generalized additive models (GAMs) in the *mgcv* package (v1.8.42; Wood 2017) in conjunction with thin-plate spline plots using the *fields* package (v14.1; Nychka, Furrer et al. 2017), following Cotter et al (2011). To account for variation in haemolymph nutrient concentrations, data were standardized using the mean (µ) and standard deviation (σ), as per Cotter et al (2011). An information theoretic approach was taken for analysis (Whittingham et al., 2006), which allows a selection of multiple candidate models to be simultaneously compared based on corrected Akaike information criteria (AICc; (Burnham & Anderson, 2004). This was carried out using the *MuMIn* package (v1.47.5; (Barton, 2023) in R which, when combined with the *mgcv* package, ranks models based on AICc values. The specific analyses varied, however they all included a ‘*Null* model’, which provided a baseline measure of variation.

## RESULTS

### Host survivorship in relation to bacterial challenge status

Overall larval mortality was 80% (n = 160/200) in insects challenged with live bacteria, 6% (n = 6/100) in those challenged with dead bacteria, and 11% (11/100) in larvae that were sham-challenged; only insects in the live-challenged group died of *X. nematophila* infection. Live-challenged larvae lived for an average of 80.1 α 104.0 h (mean α SD), whereas those in the dead-challenged and sham-challenged groups lived, on average, for another week before either dying or pupating (255.0 α 102.5 h and 235.5 α 110.8 h, respectively; **Figure 1**). Survival analysis indicated that there was a highly significant difference in the survivorships of insects in the three treatment groups overall (χ^2^ = 291.5, P < 0.0001), with the live-challenged insects suffering higher mortality rates than the two control groups (z = −8.60, P < 0.0001), which did not differ significantly in their survivorships (z = 1.33, P = 0.18).

**Figure 1.**
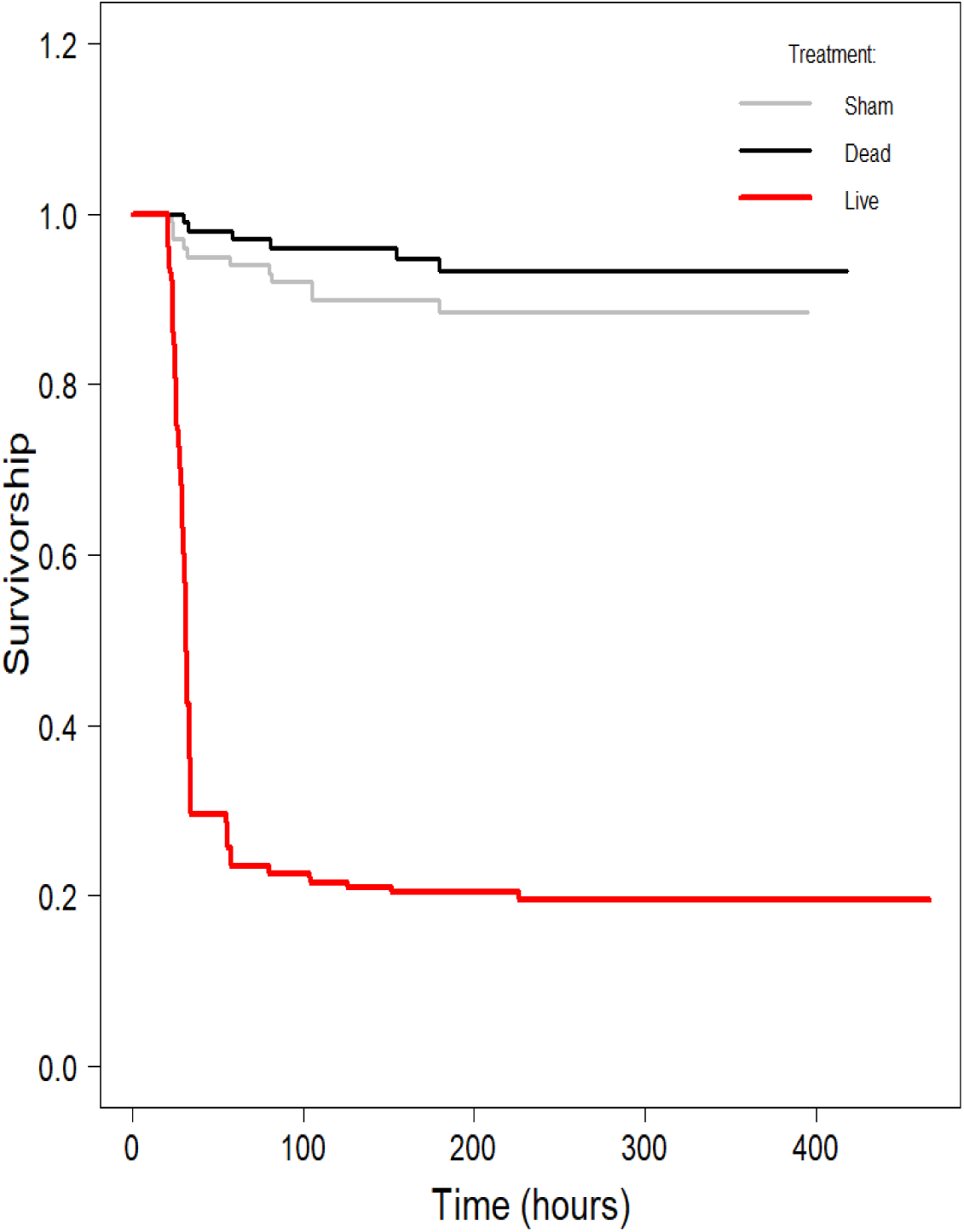
Survivorship curves (time-specific survival) for larvae in the three treatment groups. None of the larvae in the sham or dead-infected groups died of *X. nematophila* infection. Larvae were monitored until death (whether as larvae, pupae or moths), with curves censored at the timepoint where no live individuals remained.

### Host speed of death in relation to host diet

Given the very different mortality rates in the live-challenged larvae compared to those in the two control groups, the effect of diet on standardised speed of death (see Methods for calculation) was compared across these two groups separately. In the two control groups, diet had little effect on speed of death, explaining less than 5% of the variation (**Table 1A-C**), regardless of whether the larvae had been challenged with dead bacteria or sham-challenged (**Figure S2A,B**).

A previous study (Wilson, Holdbrook et al. 2020) suggested that the bacteria-challenged insects comprise three categories of individuals: those that succumb to the bacteria (and die if/when the bacterial load exceeds some critical threshold); those that successfully control the nascent bacterial population and survive infection; and those that stochastically did not receive a sufficiently large dose of cells for the bacterial population to establish. In practice, it is difficult to distinguish between these latter two categories, but we can ask whether the nutritional properties of the diet affect differently larvae that survived a live bacterial challenge (whether infected or not) versus those that did not survive (and certainly did host a growing bacterial population). This revealed that diet did not affect the speed of death of those individuals that survived the bacterial challenge (**Table 1D-F; Figure S2C**), but larvae succumbing to bacterial infection lived longer if they were fed on protein-rich diets (**Table 1D-F; Figure S2D**). Given that diet only appears to significantly affect the speed of death of larvae dying of *X. nematophila* infection, all further analyses presented here are restricted to this category of insects.

### *In vivo* bacterial growth rate in relation to host diet macronutrient composition

Bacterial counts in sampled haemolymph ~20 hours post-infection confirmed that larvae in the two control groups were free of *X. nematophila* infection. It also showed that, at the point of sampling, larvae in the live-challenge group harboured an average of around 10^4^ CFU/mL, but with most survivors (n = 30) harbouring no bacteria at sampling and with the remaining survivors averaging around 10^3^ CFU/mL (range = 5 × 10^2^ – 5 × 10^3^; n = 8). In contrast, those dying of *X. nematophila* infection averaged 10^6^ CFU/mL, with a significant number of larvae (n = 31) hosting no culturable *X. nematophila* at the point of sampling and the remainder averaging of 10^6^ CFU/mL (range = 5 × 10^2^ – 5 × 10^10^; n = 97; **Figure 2**).

**Figure 2.**
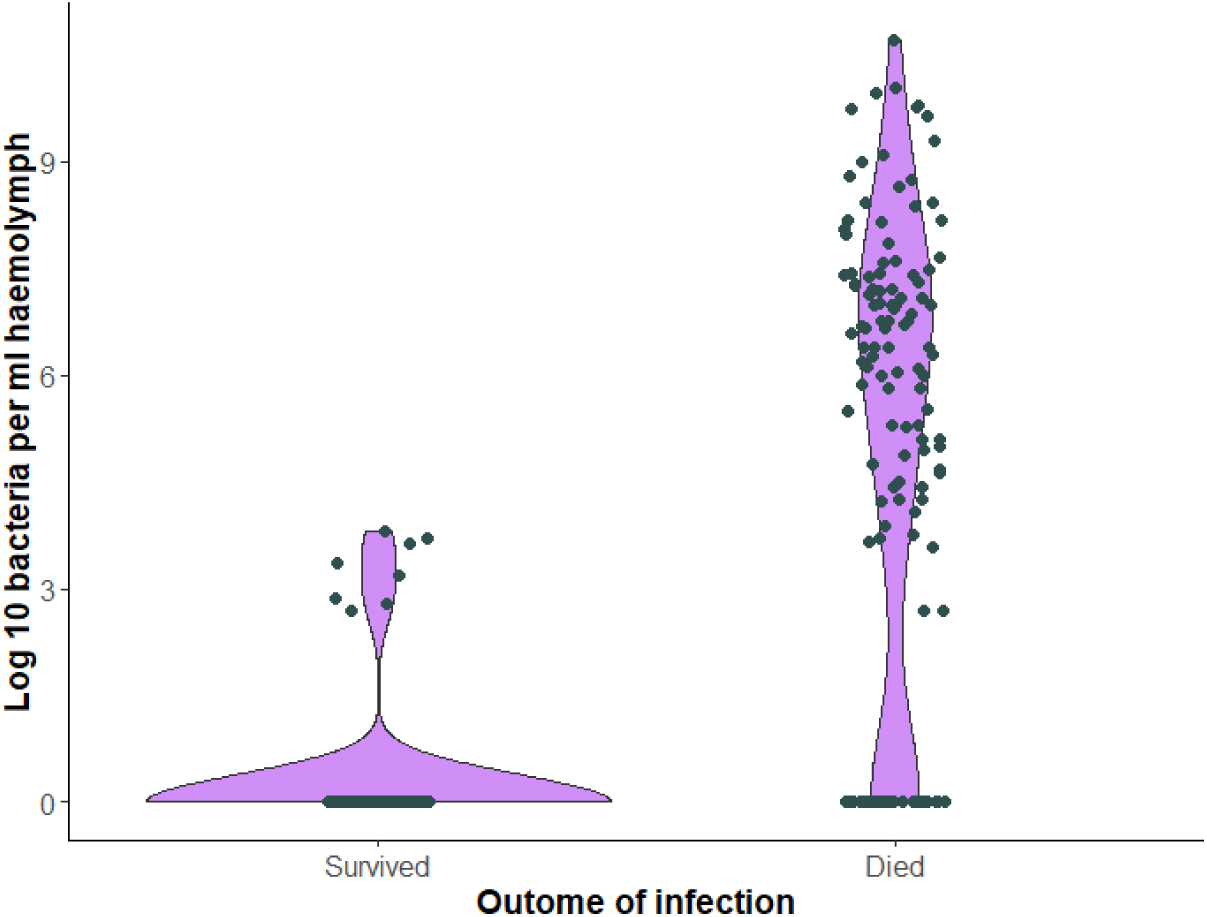
Violin plot depicting the bacterial loads of larvae at sampling with respect to whether they survived infection or died.

The bacterial load at sampling for larvae that would subsequently die of infection (i.e. the *in vivo* bacterial growth rate) was largely determined by the amount of protein in the host diet, explaining more than 30% of the variation, with little contribution from the amount of dietary carbohydrate (**Figure 3B**; **Table 2(A-C)**; bacterial load markedly decreased as the protein content of the diet increased, and increased slightly as the amount of carbohydrate increased.

**Figure 3.**
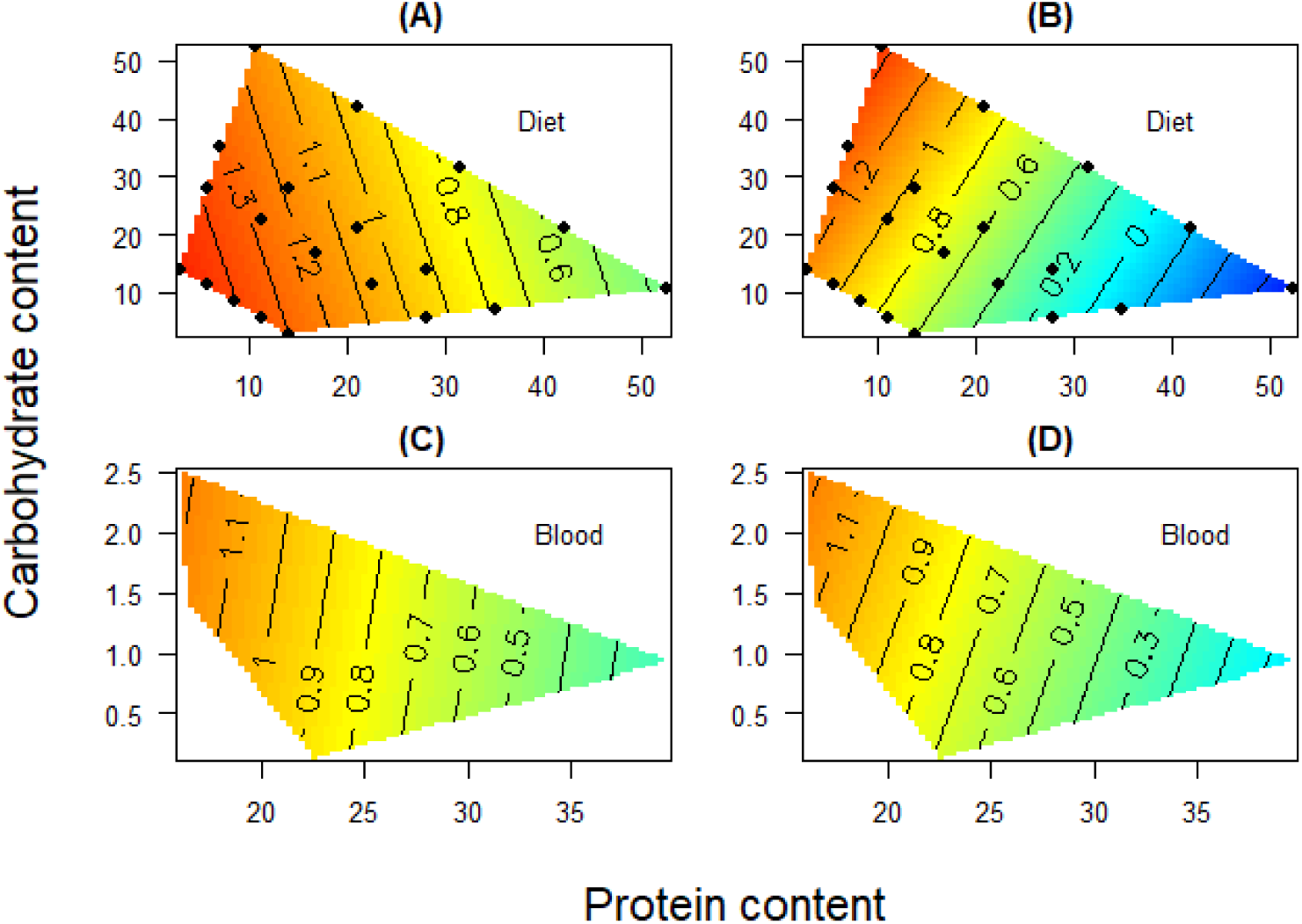
Effects of diet on standardised speed of death and bacterial growth rate in *X. nematophila*-challenged insects. Effects of protein and carbohydrate in host diet, g/100g (A,B) or haemolymph, mg/mL (C,D; see methods) on standardized (A,C) speed of death (1/lifespan, h) and (B,D) *in vivo* bacterial growth rate in larvae dying of *X. nematophila*-infection based on log(CFU/mL) and. All panels have a common z-limit.

**Table 2.** GAMs explaining standardized *in vivo* bacterial growth rate (i.e. bacterial load at sampling) in relation to the levels of protein and carbohydrate the host diet (A-C) or host haemolymph (D-H) for larvae that died of *X. nematophila*. **A,D)** Table of candidate models. df = degrees of freedom, AICc = corrected Akaike Information Criteria values; Δ AICc = difference in AICc values between the best model (lowest AICc) and the given model; w = Akaike weights; r^2^ = pseudo-r^2^ for the model. The five alternative Diet attributes listed in the first column are described in Table S1. The dependent variable in these models is log_10_(bacterial load, CFU/mL). Analysis was restricted to those larvae that died of bacterial infection. **B,E)** Parametric terms and **C,F)** Smoothed terms of the top model (model 3 - diet/Model 2 - haemolymph) and **G)** Parametric terms and **H)** Smoothed terms of the second-top model (model 4 - haemolymph).

| **A) Model** | **df** | **AIC<sub>c</sub>** | **$\Delta$ AIC<sub>c</sub>** | **w** | **$r^2$** |
| --- | --- | --- | --- | --- | --- |
| 3. Protein + Carb | 4 | 608.6 | 0.00 | 0.452 | 0.347 |
| 4. Protein * Carb | 4 | 608.6 | 0.00 | 0.452 | 0.347 |
| 1. Protein | 4 | 611.8 | 3.11 | 0.096 | 0.328 |
| 2. Carb | 4 | 653.9 | 45.28 | 0.000 | 0.074 |
| 0. No diet attributes | 2 | 659.2 | 50.59 | 0.000 | 0.000 |

| **B) Parametric terms** | **b** | **SE** | **t** | **P** |
| --- | --- | --- | --- | --- |
| intercept | 0.51206 | 0.06627 | 7.727 | < 0.0001 |

| **C) Smoothed terms** | **edf** | **Ref.df** | **F** | **P** |
| --- | --- | --- | --- | --- |
| Bleeding time | 0.00057 | 4 | 0.000 | 0.3932 |
| Protein | 0.98299 | 4 | 14.0118 | < 0.0001 |
| Carb | 1.38181 | 4 | 1.479 | 0.0153 |

| **D) Model** | **df** | **AIC<sub>c</sub>** | **$\Delta$ AIC<sub>c</sub>** | **w** | **$r^2$** |
| --- | --- | --- | --- | --- | --- |
| 2. Carb | 8 | 162.6 | 0.00 | 0.404 | 0.428 |
| 3. Protein + Carb | 10 | 163.2 | 0.61 | 0.297 | 0.447 |
| 4. Protein * Carb | 10 | 163.2 | 0.61 | 0.297 | 0.447 |
| 1. Protein | 6 | 174.0 | 11.41 | 0.001 | 0.319 |
| 0. No diet attributes | 4 | 175.9 | 13.28 | 0.001 | 0.273 |

| **E) Parametric terms** | **b** | **SE** | **t** | **P** |
| --- | --- | --- | --- | --- |
| intercept | 0.5151 | 0.0748 | 6.888 | < 0.0001 |

| **F) Smoothed terms** | **edf** | **Ref.df** | **F** | **P** |
| --- | --- | --- | --- | --- |
| Bleeding time | 3.050 | 4 | 6.67 | < 0.0001 |
| Carb | 2.763 | 4 | 4.04 | 0.0011 |

| **G) Parametric terms** | **b** | **SE** | **t** | **P** |
| --- | --- | --- | --- | --- |
| intercept | 0.5151 | 0.0748 | 6.888 | < 0.0001 |

| **H) Smoothed terms** | **edf** | **Ref.df** | **F** | **P** |
| --- | --- | --- | --- | --- |
| Bleeding time | 3.419 | 4 | 6.604 | <0.0001 |
| Protein | 1.487 | 4 | 1.261 | 0.025 |
| Carb | 2.273 | 4 | 2.816 | 0.003 |

### *In vivo* bacterial growth rate and speed of death in relation to haemolymph macronutrient composition

In a previous paper, we explored the effects of host diet on the nutritional composition of the host haemolymph – the environment in which *X. nematophila* would grow (Holdbrook, Andongma, et al., 2024). To establish how the nutritional composition of *S. littoralis* haemolymph translates into bacterial growth rates and host speed of death of larvae challenged with live *X. nematophila*, we quantified the relationships between these traits in the current experiment with the average haemolymph composition of larvae fed on each of the twenty diets (Holdbrook, Andongma, et al., 2024). It should be noted that both haemolymph protein and haemolymph carbohydrate are positively correlated with the relative amount of protein and carbohydrate in the host diet, respectively: *r* (protein) = 0.685, df = 115, P < 0.0001; *r* (carbohydrate) = 0.739, df = 115, P < 0.0001). It is also pertinent to note that, in the host haemolymph, these two macronutrients are strongly negatively correlated (*r* = −0.519, df = 115, P < 0.0001), making it difficult to distinguish their independent effects.

In terms of *in vivo* bacterial growth rate, as the putative amount of haemolymph carbohydrate increased, so too did the bacterial load of dying larvae (**Figure 3D**, **Table 2D-F**, R^2^ = 0.428). Given that dietary protein has previously been implicated in *X. nematophila* growth rates, its higher R^2^ (0.447) and that the heatmap (**Figure 3D**) appears to suggest a strong protein effect, we also present the outputs from the second-best model which includes haemolymph protein and its haemolymph carbohydrate for comparison (**Table 2G,H**). This suggests that the *in vivo* bacterial growth rate is associated with a strong interaction between the putative amount of protein and carbohydrate in the host haemolymph, with *in vivo X. nematophila* growth increasing with haemolymph carbohydrate and decreasing with haemolymph protein.

The speed of death of larvae dying of *X. nematophila* infection was largely determined by the putative amount of protein in the larval haemolymph, with all three of the top models that include protein explaining 36 - 46% of the variation (**Figure 3C**; **Table 3**); in contrast, carbohydrate alone explained little variation in the speed of death.

**Table 3.**
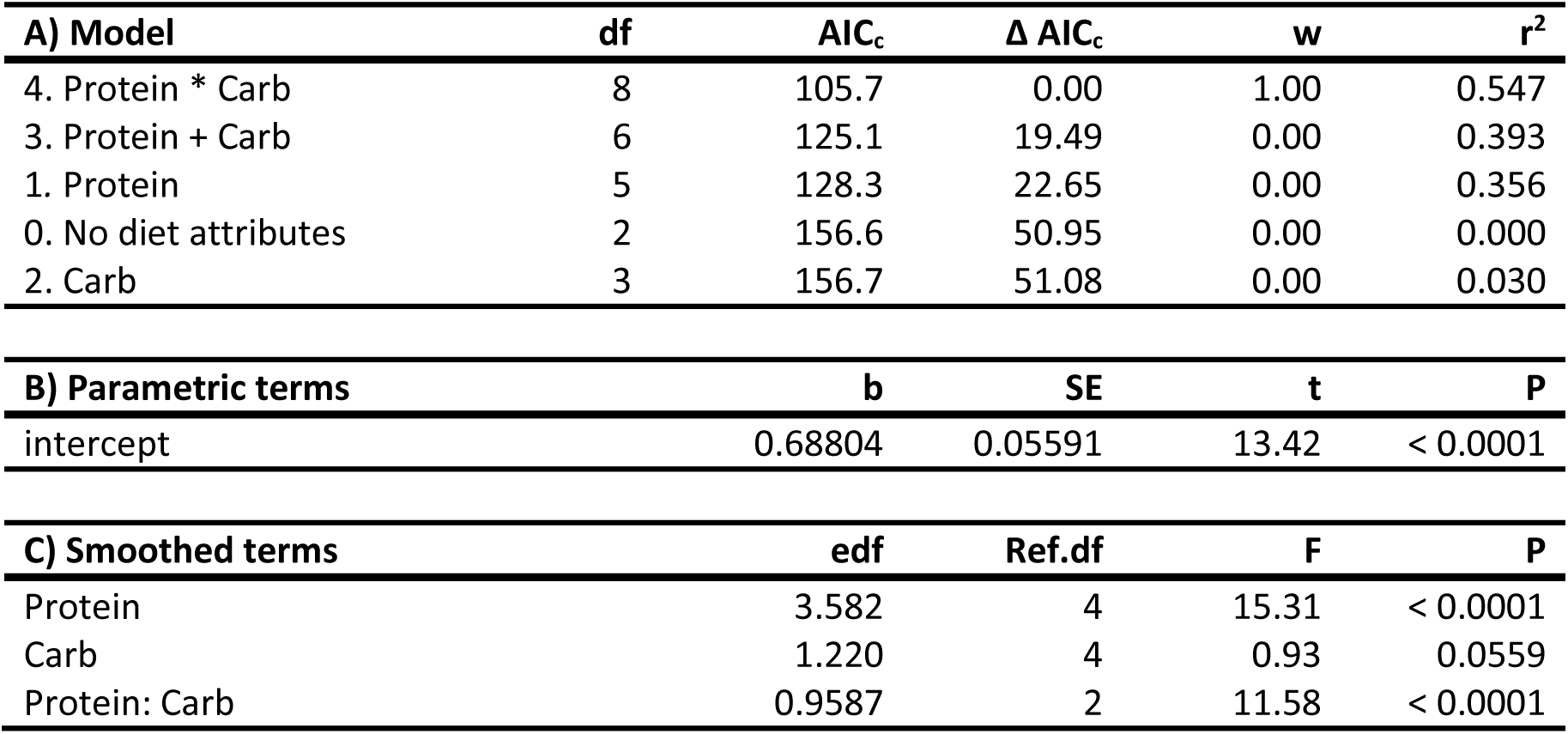
GAMs explaining the standardised speed of death of larvae dying of *X. nematophila* infection in relation to the putative levels of protein and carbohydrate in the host haemolymph pre-infection. **A)** Table of candidate models. df = degrees of freedom, AICc = corrected Akaike Information Criteria values; Δ AICc = difference in AICc values between the best model (lowest AICc) and the given model; w = Akaike weights; r^2^ = pseudo-r^2^ for the model. The dependent variable in these models is speed of death (1/h). **B)** Parametric terms and **C)** Smoothed terms of the top model (model 2).

### *In vitro* bacterial growth rate in relation to putative host diet based on synthetic haemolymphs – ‘nutribloods’

When *in vitro* bacterial growth rate (as measured spectrophotometrically by maximum OD at 600 nm, *maxOD*) was analysed in relation to the amount of protein and carbohydrate in the nutribloods (and by extension in the host’s haemolymph), this revealed that as the protein content of the haemolymph increased, so the *in vitro* growth rate of *X. nematophila* declined (**Figure 4A**; **Table 4A-C**). It should be noted, however, that all four diet models explained a similar amount of variation in the bacterial growth rate (*r*^2^ = 0.210 – 0.258), with the greatest amount of variation explained by a model that included both the amount of protein and the amount of carbohydrate in the nutriblood. This is due to the fact that, the amount of protein and carbohydrate in the synthetic haemolymphs are strongly negatively correlated (Pearson’s *r* = −0.751, df = 18, P = 0.0001).

**Figure 4.**
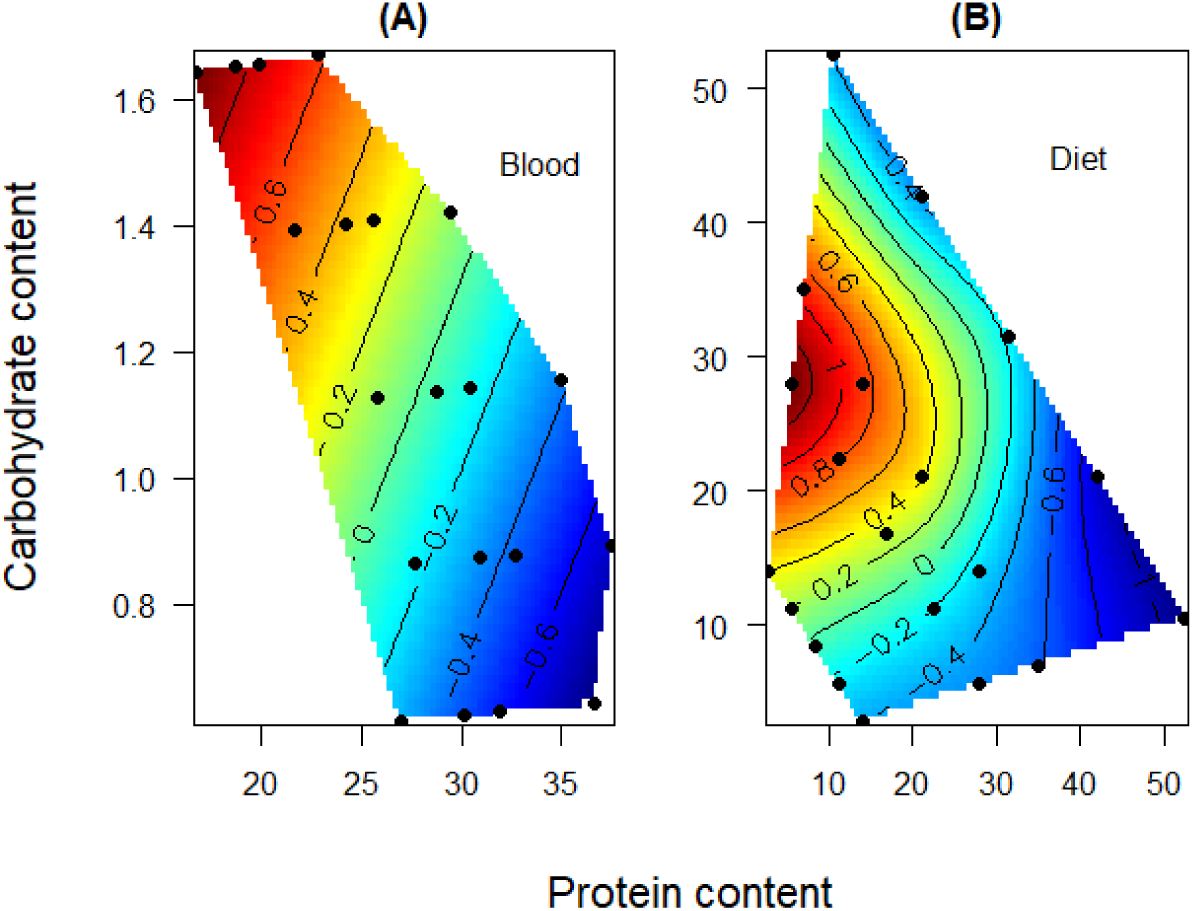
The effects of nutrients on *in vitro* bacterial growth rates. Relationships between (A) nutriblood macronutrients, mg/ml (B) putative host diet, g/100g, on standardized *in vitro* bacterial growth rate of *X. nematophila*, as measured by log maximum OD at 600 nm.

**Table 4.** GAMs explaining the in vitro growth rate of X. nematophila in relation to the levels of protein and carbohydrate in (A-C) the synthetic haemolymphs (‘nutribloods’) or (D-F) the host diet. **A,D)** Table of candidate models. df = degrees of freedom, AICc = corrected Akaike Information Criteria values; Δ AICc = difference in AICc values between the best model (lowest AICc) and the given model; w = Akaike weights; r2 = pseudo-r2 for the model. The five alternative Diet attributes listed in the first column are described in Table S1. The dependent variable in these models is the maximum OD at 600 nm. **B,E)** Parametric terms and **C,F)** Smoothed terms of the top model (model 1 – nutribloods, model 3 - diets).

| **A) Model** | **df** | **AIC<sub>c</sub>** | **$\Delta$ AIC<sub>c</sub>** | **w** | **r<sup>2</sup>** |
| --- | --- | --- | --- | --- | --- |
| 1. Protein | 2 | -196.8 | 0.00 | 0.296 | 0.228 |
| 3. Protein + Carb | 4 | -196.7 | 0.15 | 0.275 | 0.258 |
| 4. Protein * Carb | 4 | -196.7 | 0.15 | 0.275 | 0.258 |
| 2. Carb | 2 | -195.5 | 1.32 | 0.153 | 0.210 |
| 0. No diet attributes | 1 | -182.8 | 14.01 | 0.000 | 0.000 |

| **B) Parametric terms** | **b** | **SE** | **t** | **P** |
| --- | --- | --- | --- | --- |
| intercept | 0.4124 | 0.0057 | 72.07 | < 0.0001 |

| **C) Smoothed terms** | **edf** | **Ref.df** | **F** | **P** |
| --- | --- | --- | --- | --- |
| Protein | 0.4124 | 0.0046 | 89.58 | < 0.0001 |

| **D) Model** | **df** | **AIC<sub>c</sub>** | **$\Delta$ AIC<sub>c</sub>** | **w** | **r<sup>2</sup>** |
| --- | --- | --- | --- | --- | --- |
| 3. Protein + Carb | 10 | -211.1 | 0.00 | 0.999 | 0.500 |
| 2. Carb | 7 | -197.7 | 13.41 | 0.001 | 0.323 |
| 4. Protein * Carb | 3 | -193.9 | 17.18 | 0.000 | 0.206 |
| 1. Protein | 3 | -193.7 | 17.41 | 0.000 | 0.198 |
| 0. No diet attributes | 1 | -182.8 | 28.33 | 0.000 | 0.000 |

| **E) Parametric terms** | **b** | **SE** | **t** | **P** |
| --- | --- | --- | --- | --- |
| intercept | 1.89370 | 0.07760 | 24.40 | < 0.0001 |

| **F) Smoothed terms** | **edf** | **Ref.df** | **F** | **P** |
| --- | --- | --- | --- | --- |
| Protein | 3.451 | 9 | 2.599 | < 0.0001 |
| Carb | 5.091 | 9 | 3.874 | < 0.0001 |

When these same (max OD_600_) data were analysed in terms of the putative amount of protein and carbohydrate in the host diet, as back calculated from the nutribloods (**Figure 4B**; **Table 4D-F**), *in vitro* bacterial growth rate was highest on putative low-protein / medium-carbohydrate diets.

### *In vivo* and *in vitro* bacterial growth rate in relation to host diet

Finally, to determine how much variation in bacterial growth rate *in vivo* differs from that of bacteria growing in the synthetic haemolymphs, we produced a dataset that included both the standardised *in vivo* growth rates for larvae fed the twenty chemically-defined diets and the standardised *in vitro* growth rates for bacteria grown in the twenty nutribloods mimicking the average nutritional properties of haemolymph collected from larvae feeding on the same twenty diets. We then asked how much of the relationship between bacterial growth rate and the nutritional properties of host ‘diet’ depended on whether the bacteria were growing *in vivo* or *in vitro*.

To do this, we included a dummy *Treatment* term classifying bacterial growth rate as *in vivo* or *in vitro* and added this as a factor in the standard GAMs, as previously, as well as in comparable models in which diet was allowed to interact with *Treatment*. This revealed that the top four models included both protein and carbohydrate as significant nutritional terms (*r*^2^_models 3a, 3b, 4a, 4b_ = 0.319 - 0.358), and that the interaction with *Treatment* explained around 4% additional variation in standardised bacterial growth rates than models that included *Treatment* only as an additive parametric term (i.e. *r*^2^ = ~0.36 *versus* ~0.32) (**Table 5**). This was due to the fact that the interaction between dietary protein and carbohydrate was much more important in the bacteria that were growing *in vitro* than *in vivo* (**Figure 4**), as reflected in the more non-monotonic contours with respect to the carbohydrate axis of the former.

**Table 5.** GAMs explaining variation in the *in vivo* and *in vitro* growth rate of X. nematophila in relation to the levels of protein and carbohydrate in the host diet. **A)** Table of candidate models. df = degrees of freedom, AICc = corrected Akaike Information Criteria values; Δ AICc = difference in AICc values between the best model (lowest AICc) and the given model; w = Akaike weights; r2 = pseudo-r2 for the model. The five alternative haemolymph attributes listed in the first column are described in Table 1. The dependent variable in these models is the standardized bacterial growth rate (in vivo = log10-bacterial load at sampling for individuals dying of X. nematophila infection and in vitro = max OD at 600 nm). All models also included Treatment (i.e. *in vivo* or *in vitro*) as a parametric term in the model. **B)** Parametric terms and **C)** Smoothed terms of the top model (model 4b).

| **A) Model** | **df** | **AIC<sub>c</sub>** | **$\Delta$ AIC<sub>c</sub>** | **w** | **r<sup>2</sup>** |
| --- | --- | --- | --- | --- | --- |
| 4b. Protein * Carb / Treatment | 8 | 456.4 | 0.00 | 0.501 | 0.356 |
| 3b. Protein + Carb / Treatment | 9 | 456.5 | 0.15 | 0.466 | 0.358 |
| 4a. Protein * Carb | 6 | 463.3 | 6.92 | 0.016 | 0.319 |
| 3a. Protein + Carb | 6 | 463.3 | 6.92 | 0.016 | 0.319 |
| 1a. Protein | 4 | 470.1 | 13.71 | 0.001 | 0.283 |
| 1b. Protein / Treatment | 5 | 472.1 | 15.76 | 0.000 | 0.282 |
| 2b. Carb / Treatment | 7 | 517.9 | 61.56 | 0.000 | 0.095 |
| 2a. Carb | 4 | 522.1 | 65.72 | 0.000 | 0.059 |
| 0a. No diet attributes | 3 | 532.0 | 75.58 | 0.000 | 0.000 |
| 0b. No diet attributes / Treatment | 3 | 532.0 | 75.58 | 0.000 | 0.000 |

| **B) Parametric terms** | **b** | **SE** | **t** | **P** |
| --- | --- | --- | --- | --- |
| intercept | -0.04087 | 0.10504 | -0.389 | 0.698 |
| Treatment (*in vivo*) | 0.07691 | 0.12688 | 0.606 | 0.545 |

| **C) Smoothed terms** | **edf** | **Ref.df** | **F** | **P** |
| --- | --- | --- | --- | --- |
| Protein / *in vitro* | 0.855 | 4 | 1.473 | 0.0088 |
| Protein / *in vivo* | 0.994 | 4 | 14.299 | < 0.0001 |
| Carb / *in vitro* | 0.000 | 4 | 0.000 | 0.7212 |
| Carb / *in vivo* | 1.359 | 4 | 1.354 | 0.0198 |
| Protein : Carb / *in vitro* | 1.789 | 2 | 8.678 | < 0.0001 |
| Protein : Carb / *in vivo* | 0.000 | 2 | 0.000 | 0.4975 |

### Correlation between bacterial growth *in vivo* and *in vitro*

Finally, since we have made comparisons in bacterial growth *in vivo* and *in vitro* using two different methods (CFU/mL and OD_600_, respectively), we compared the relationship between the two. Across the 20 diets and nutribloods, the *in vivo* and *in vitro* bacterial growth rates were significantly positively correlated with each other (Pearson’s *r* = 0.251, df = 203, P = 0.0003), suggesting that the synthetic haemolymphs provide a reasonable reflection of the nutritional properties of real haemolymph and that the two methods for quantifying bacterial growth can be used fairly interchangeably. Indeed, these findings concur with other studies that have compared bacterial growth using both of these methods (González-Pérez et al., 2019; Karamba & Ahmad, 2019; Kim et al., 2012; Kowalski et al., 1998).

## DISCUSSION

This study aimed to combine data from *in vitro* and *in vivo* analyses of bacterial performance on 20 chemically-defined diets to determine the relative importance of top-down (immunological) from bottom-up (resource) effects of host diet using as a model system a generalist caterpillar host and its extra-cellular bacterial pathogen. The study revealed that host diet, most notably intake of dietary protein, markedly affected the rate at which the insects died of the bacterial infection, which in turn was determined by the effects of diet on the rate of *in vivo* bacterial growth, with hosts dying when the bacterial population had reached a critical threshold level (Wilson et al., 2020). This was largely driven by, or at least was strongly correlated with, the putative amount of protein in the insect host’s haemolymph (blood). Using detailed information on the nutritional composition of haemolymph from caterpillars feeding on each of the 20 diets (Holdbrook, Andongma, et al., 2024), we generated 20 synthetic haemolymphs (nutribloods) mimicking the nutritional compositions of caterpillars feeding on each of the 20 diets (Holdbrook, Randall, et al., 2024) and quantified relative bacterial performance *in vitro* in the absence of the host’s immune system. This revealed a similar pattern to that seen *in vivo*, with bacterial growth rate being highest in nutribloods low in proteins and high in carbohydrates. A statistical analysis comparing the bacterial performance profiles *in vivo* and *in vitro* revealed only marginal differences between the two, consistent with the bacterial population being limited mainly by the direct effects of nutrients on bacterial growth rate (bottom-up effects), rather than by the indirect effects of nutrients on its host’s immune responses (top-down effects).

### *In vivo* bacterial performance

When larvae were sampled for bacteria during the exponential phase of bacterial growth in the host, the concentration of *X. nematophila* cells in the haemolymph was strongly negatively correlated with the amount of protein in the host diet and weakly positively correlated with the amount of dietary carbohydrate (Figure 3). This is, in part, consistent with our previous study using just six chemically-defined diets (Wilson et al., 2020). However, using 20 diets in this study allowed us to also detect the positive effect of dietary carbohydrate on bacterial performance, which was not observed in the 6 diets experiment (Wilson et al., 2020), and suggesting that carbohydrates or their metabolites are being used by the bacteria as a food source. We also observed that the speed at which larvae succumbed to *X. nematophila* infection was strongly negatively related to the amount of protein in the host diet, and putative levels of protein in the larval haemolymph (Figure 4), presumably because the bacterial population was growing at a slower rate when the larvae ate protein-rich diets and had high levels of protein in the haemolymph (Holdbrook, Andongma, et al., 2024; Wilson et al., 2020).

This observed ‘protein effect’ could be due to a negative impact of the host’s immune response on the *in vivo* bacterial growth rate. Consistent with this, a previous study using this system revealed that dietary protein facilitates the functional immune responses of *S. littoralis* against a *X. nematophila* challenge, though the effect was relatively small, explaining less than 12% of the variation in the immune response (Cotter et al., 2019). Disentangling these top-down immune responses from bottom-up resource utilization effects is challenging in host-parasite interactions but has been attempted in a number of systems (Griffiths et al., 2015; Haydon et al., 2003; Metcalf et al., 2011; Mideo & Reece, 2012; Moore et al., 2018; Ramiro et al., 2016) providing evidence for both top-down and bottom-up effects. For example, Metcalf et al (2011) used a parameterised mathematical model to distinguish these effects in a mouse-malaria host-parasite system. However, such an approach requires fine-resolution time-series data on the relative proportions of infected and non-infected cells and is not appropriate for an extra-cellular parasite like *X. nematophila*. A similar approach was taken by Moore et al (2018) to explore the viral dynamics in humans vaccinated with the attenuated live virus that causes yellow fever. As with the mouse-malaria model, however, these infections are long-lasting and require repeated sampling of individuals over time, which is logistically challenging for short-lived infections in insects.

An alternative approach to establishing the relative importance of top-down immune regulation and bottom-up resource utilization is to remove the effects of the host immune response by quantifying *in vitro* parasite growth dynamics. Although previous studies have compared the effects of different nutrients on *in vitro* microbial growth, these usually involve batch cultures with generic media containing multiple nutrients that are simultaneously varied or the modification of a single dietary component (Bowen et al., 2012; Kooliyottil et al., 2014; Pulkkinen et al., 2018). In contrast, in the present study we systematically varied key nutrients in twenty solutions that mimicked, as far as possible, the nutritional conditions that the bacterium could expect to face within its host following different dietary regimes (Holdbrook, Andongma, et al., 2024).

### *In vitro* bacterial performance

For *S. littoralis* fed on the 20 chemically-defined diets, the haemolymph protein and carbohydrate levels are strongly negatively correlated with each other, making it difficult to distinguish their independent effects. However, the *in vitro* bacterial growth experiment clearly shows that bacteria grow best in nutribloods rich in carbohydrates and low in proteins (Figure 4A). When these nutribloods are back-transformed to the host diets that they reflect, it becomes clear that *in vitro* bacterial growth peaks on diets with intermediate levels of carbohydrates and low levels of protein (Figure 4B).

When we compared this pattern of *in vitro* growth with that observed *in vivo*, the patterns were broadly similar, with both exhibiting a strong negative ‘protein effect’ (*cf.* Figure 4A,B). Moreover, when the standardized growth patterns were compared statistically, there was no overall significant difference between the two, with the model that distinguished between *in vivo* and *in vitro* growth explaining less than 5% more variation in bacterial growth rate than the model without. This was a consequence of *in vitro* growth exhibiting a stronger interaction between dietary carbohydrate and protein. This finding is consistent with bacterial growth largely being regulated by bottom-up effects of the haemolymph nutrients, especially haemolymph protein, rather than top-down effects of host nutrition on its immune response, as reflected in their relatively weak functional immune response to *X. nematophila* infection (Cotter et al., 2019). A number of studies have used *in vitro* studies of bacteria to predict growth properties *in vivo*, for example in the screening of potential probiotic bacteria (Foligne et al., 2007; Grangette et al., 2005) or potential antibiotics (Ono et al., 1996; Schmidtchen & Puthia, 2022). However, as far as we are aware, this is the first time that *in vitro* and *in vivo* approaches have been combined to study the effects of nutrition on a host-parasite interaction.

### Proteins may interfere with osmoregulation

Abisgold & Simpson (1987) found that increasing protein concentration in the diet of *Locusta migratoria* increased haemolymph amino acid concentration, which in turn raised haemolymph osmolality. Osmoregulation, or a cell’s ability to adapt to changes in their osmotic environment, is important for the maintenance of turgor pressure across the cellular membrane (Csonka & Hanson, 1991; Kempf & Bremer, 1998). The osmoregulatory ability of a cell, in turn, determines its ability to counteract osmotic stress and therefore its ability to proliferate (Csonka, 1989; Tempest et al., 1970). The findings of Abisgold & Simpson (1987) highlight changes in osmolality as a possible mechanism for the observed ‘protein effect’. Indeed, Wilson et al (2020) observed that the osmolality of *S. littoralis* haemolymph increased as the amount of protein in the host diet increased. Moreover, both *in vitro* and *in vivo* studies showed that *X. nematophila* growth rate declines with increasing osmolality, providing a potential mechanism for the observed negative effect of host dietary protein on bacterial growth and its positive effect on host survival (see Figure 5 in Wilson et al (2020)) It is pertinent to note that many animals self-medicate in response to infection (e.g. Erler et al., 2024). For example, the congener, *Spodoptera exempta* caterpillars increase their intake of dietary protein in response to bacterial infection (Povey et al., 2009) consistent with them triggering this negative ‘protein effect’ observed in this study.

## Conclusion

The aim of this study was to use an *in vitro* system to determine whether the host nutritional effects on the pathogen observed could be due to direct (bottom-up) effects in addition to the previously observed host-mediated (top-down) immunological effects (Cotter et al., 2011, 2019). We provide strong evidence that bacterial growth is primarily regulated by the bottom-up effects of host nutrition, particularly via the negative effects of haemolymph protein, most likely via inducing osmotic stress on the bacterial cells (Wilson et al., 2020). Moreover, via the creation of synthetic insect haemolymphs (Holdbrook, Randall, et al., 2024), we provide a tractable experimental framework for testing the role of nutrition in host-pathogen and host-commensal relationships in insect blood. For example, one potential use of this system is to elucidate the nutritional requirements of the nematode symbionts, such as *S. carpocapsae* that remain unknown (Richards & Goodrich-Blair, 2009).

