## Supplementary material for "Combining *in vivo* and *in vitro* approaches to better understand host-pathogen interactions"

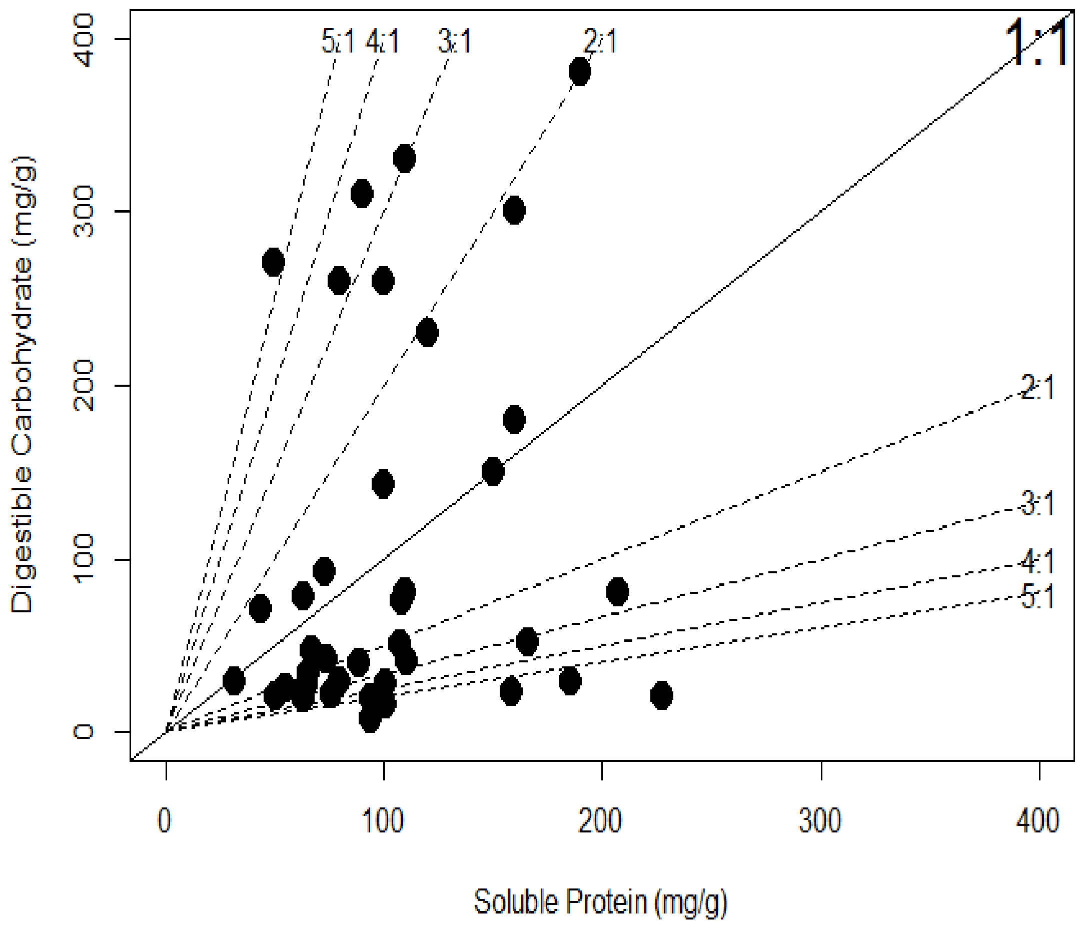


**Figure S1. The macronutrient composition of plants typically fed on by the generalist caterpillar, *S. littoralis*.** The data were taken from Scott Brown, Simmonds and Blaney (2002) and Wilson, Ruiz and Davidowitz (2019).

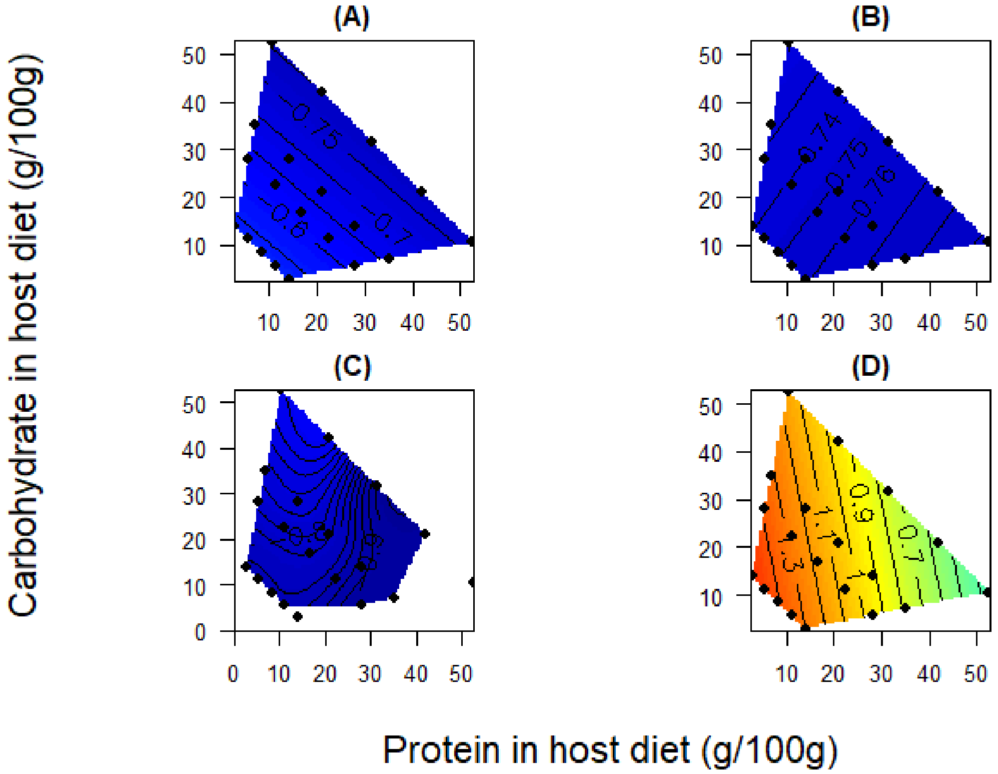


**Figure S2. Effects of diet on standardised speed of death in *X. nematophila*-challenged insects.** (A) Dead-infected, injected with heat-killed bacteria in nutrient broth; (B) sham-infected, injected with nutrient broth only; (C) Live-infected – survived, injected with live bacteria and survived; (D) Live-infected – died, injected with live bacteria and died. The z-value range in this plot is fixed such that low values appear dark blue and high values are increasingly warmer colours.

**Table S1: Twenty diets fed to *Spodoptera littoralis* caterpillars varying in their ratios and concentrations of protein and carbohydrate.** For full details of the diets used see Table S1 in Cotter et al (2019)**.**

| Diet | P:C ratio | Diet concentration (g/100g) | % protein | Protein (g/100g) | Carbohydrates (g/100g) |
|---|---|---|---|---|---|
| 1 | 1:5 | 63 | 17 | 10.5 | 52.5 |
| 2 | 1:5 | 42 | 17 | 7 | 35 |
| 3 | 1:5 | 33.6 | 17 | 5.6 | 28 |
| 4 | 1:5 | 16.8 | 17 | 2.8 | 14 |
| 5 | 1:2 | 63 | 33 | 21 | 42 |
| 6 | 1:2 | 42 | 33 | 14 | 28 |
| 7 | 1:2 | 33.6 | 33 | 11.2 | 22.4 |
| 8 | 1:2 | 16.8 | 33 | 5.6 | 11.2 |
| 9 | 1:1 | 63 | 50 | 31.5 | 31.5 |
| 10 | 1:1 | 42 | 50 | 21 | 21 |
| 11 | 1:1 | 33.6 | 50 | 16.8 | 16.8 |
| 12 | 1:1 | 16.8 | 50 | 8.4 | 8.4 |
| 13 | 2:1 | 63 | 67 | 42 | 21 |
| 14 | 2:1 | 42 | 67 | 28 | 14 |
| 15 | 2:1 | 33.6 | 67 | 22.4 | 11.2 |
| 16 | 2:1 | 16.8 | 67 | 11.2 | 5.6 |
| 17 | 5:1 | 63 | 83 | 52.5 | 10.5 |
| 18 | 5:1 | 42 | 83 | 35 | 7 |
| 19 | 5:1 | 33.6 | 83 | 28 | 5.6 |
| 20 | 5:1 | 16.8 | 83 | 14 | 2.8 |

Cotter, Sheena C., Catherine E. Reavey, Yamini Tummala, Joanna L. Randall, Robert Holdbrook, Fleur Ponton, Stephen J. Simpson, Judith A. Smith, and Kenneth Wilson. 2019. "Diet Modulates the Relationship between Immune Gene Expression and Functional Immune Responses." *Insect Biochemistry and Molecular Biology* 109 (June): 128–41.

Scott Brown, Alison S., Monique S. J. Simmonds, and Walter M. Blaney. 2002. "Relationship between Nutritional Composition of Plant Species and Infestation Levels of Thrips." *Journal of Chemical Ecology* 28 (12): 2399–2409.

Wilson, J. Keaton, L. Ruiz, and G. Davidowitz. 2019. "Dietary Protein and Carbohydrates Affect Immune Function and Performance in a Specialist Herbivore Insect (Manduca Sexta)." *Physiological and Biochemical Zoology: PBZ* 92 (1): 58–70.
